# Hyperactive intestinal proteolysis underlies *smn-1* mutant phenotypes

**DOI:** 10.64898/2026.08.11.744262

**Authors:** Anusha Iyengar, Lindsey Philips, Adam Norris

## Abstract

Many neurological diseases are caused by mutations in broadly-expressed genes, but the basis for their neuron-specific manifestation is unclear. In Spinal Muscular Atrophy (SMA), loss of the ubiquitously-expressed spliceosome assembly factor *SMN1* causes selective degeneration of motor neurons, leading to progressive neuromuscular decline. We explored the mechanisms of this cell-specific vulnerability using SMA models in the nematode *C. elegans*, which likewise exhibit progressive neuromuscular defects upon loss of *smn-1*. Surprisingly, our results show that the intestine – not neurons or muscle – is the selectively-vulnerable tissue causing *smn-1* phenotypes. RNA-Seq reveals that loss of intestinal *smn-1* causes specific global splicing defects, accompanied by robust transcriptional activation of the Intracellular Pathogen Response (IPR), a stress pathway enriched for ubiquitin-proteostasis genes. Consistent with this, *smn-1* mutants exhibit elevated levels of proteasome activity. Pharmacological proteasome inhibition rescues many of the *smn-1* mutant defects, as does deletion of specific components of the IPR pathway. These results reveal how the ubiquitously-expressed SMN-1 protein is required in a single tissue to avoid degenerative defects caused by hyperactive proteasome activity, contributing to our understanding of how mutations in ubiquitously-expressed genes can cause highly cell-specific pathologies.

**SIGNIFICANCE STATEMENT:** Many ubiquitously expressed genes cause highly selective neurodegenerative diseases, such as Huntington’s disease and Amyotrophic Lateral Sclerosis. The basis for this cell-specific vulnerability remains unclear. We address this question for *smn-1* in *C. elegans*. We show that *smn-1* is indeed required in a cell-specific manner, but unexpectedly not in neurons, but rather in the intestine. Both survival defects and behavioral phenotypes originate from intestinal loss of *smn-1*. We show that these defects are caused by hyperactive protein degradation and immune responses, and that mutant defects can be resolved by reducing these proteostasis and immune pathways using genetics or pharmacology. These results shed light on how a single tissue/cell can dictate the effects of a systemic genetic disease.

## INTRODUCTION

Mutations in *SMN1* cause Spinal Muscular Atrophy (SMA), a neuromuscular disease characterized by degeneration of spinal cord motor neurons, progressive muscle wasting and death^1,2^. In most organisms, the survival motor neuron (SMN) protein is required for survival beyond early developmental stages. For example, loss of SMN in mice or zebrafish results in embryonic lethality^3^. In humans, however, a genomic duplication event led to the generation of a second *SMN* gene, *SMN2*^4,5^, which produces partially-functional SMN^6^. As a result, *SMN1* mutations in humans are often not embryonically lethal, allowing survival for months to years before the onset of motor neuron degeneration^1^.

The SMN protein is required for assembly of spliceosomal snRNP complexes, and is expressed ubiquitously across all cell types^7–9^. Why the loss of a ubiquitously-expressed gene causes motor neuron–specific pathology remains a fundamental question in the field^3^. While alterations in various tissues and cell types have been observed in *SMN1* mutants^3,10–12^, motor neuron degeneration is the dominant phenotypic driver and cause of death^13^. Understanding the cell-specific pathogenesis and non-cell-autonomous effects across tissues is broadly relevant, since several other ubiquitously expressed genes also cause neuron-specific genetic pathologies in humans, *e.g.* in Huntington’s Disease and Amyotrophic Lateral Sclerosis^14,15^.

We set out to genetically interrogate the tissue- and cell-specific requirement for *smn-1* in *C. elegans*, which is particularly well-suited for testing tissue specificity and non-cell-autonomous tissue crosstalk *in vivo*. As in other species, *smn-1* null mutants die before reaching adulthood and exhibit progressive neuromuscular defects during larval development. These defects are presumed to be caused by loss of *smn-1* in motor neurons and/or muscle^16–18^; however, the tissue-specific requirement for *smn-1* has not been systematically tested. To address this, we used the FLP/FRT recombination system to generate knockouts (KOs) of *smn-1* in specific tissues, and in subsets of neurons, to identify the tissue-specific requirements for *smn-1*.

Surprisingly, we found that neither neuronal nor muscle KOs reproduce *smn-1* mutant phenotypes. In contrast, intestinal KOs closely phenocopy the null mutant, while intestinal rescue of *smn-1* is sufficient to rescue both behavioral and survival defects. Transcriptomic analysis reveals specific global splicing defects in *smn-1* KOs, as well as robust upregulation of immune and proteostasis genes characteristic of the Intracellular Pathogen Response (IPR). Consistent with these molecular changes, *smn-1* mutants share phenotypic hallmarks of the IPR, including stress resistance at the expense of shortened lifespan. *smn-1* mutants are partially rescued by either pharmacological proteasome inhibition or genetic deletion of specific IPR gene components. Thus, in worms the intestine rather than the neuromuscular system is particularly vulnerable to loss of *smn-1*, due in part to its particular role in pathogen sensing and proteostasis.

## RESULTS

### Tissue-specific *smn-1* deletions using the FLP/FRT system

To investigate the tissue- and cell-specific requirements for *smn-1* in *C. elegans*, we first visualized the endogenous expression of the SMN-1 protein. Using CRISPR/Cas9, we tagged the endogenous *smn-1* locus with a C-terminal mScarlet translational fusion, and observed widespread expression across tissues and cell types (Figure 1A). Expression is particularly high in the germline (Figure 1A), consistent with prior transgenic reporters and the known importance of maternally supplied *smn-1* mRNA during early embryogenesis^19^. To further resolve SMN-1 expression in specific somatic tissues, we co-labeled with tissue-specific GFP reporters, and found robust expression of SMN-1 in all cells we tested, including neurons, muscle, and intestine (Figure 1B). This is consistent with the ubiquitous expression previously reported for *smn-1* transgenes in worms^19, 20^ and in most other organisms^4,21–24^.

**Figure 1:**
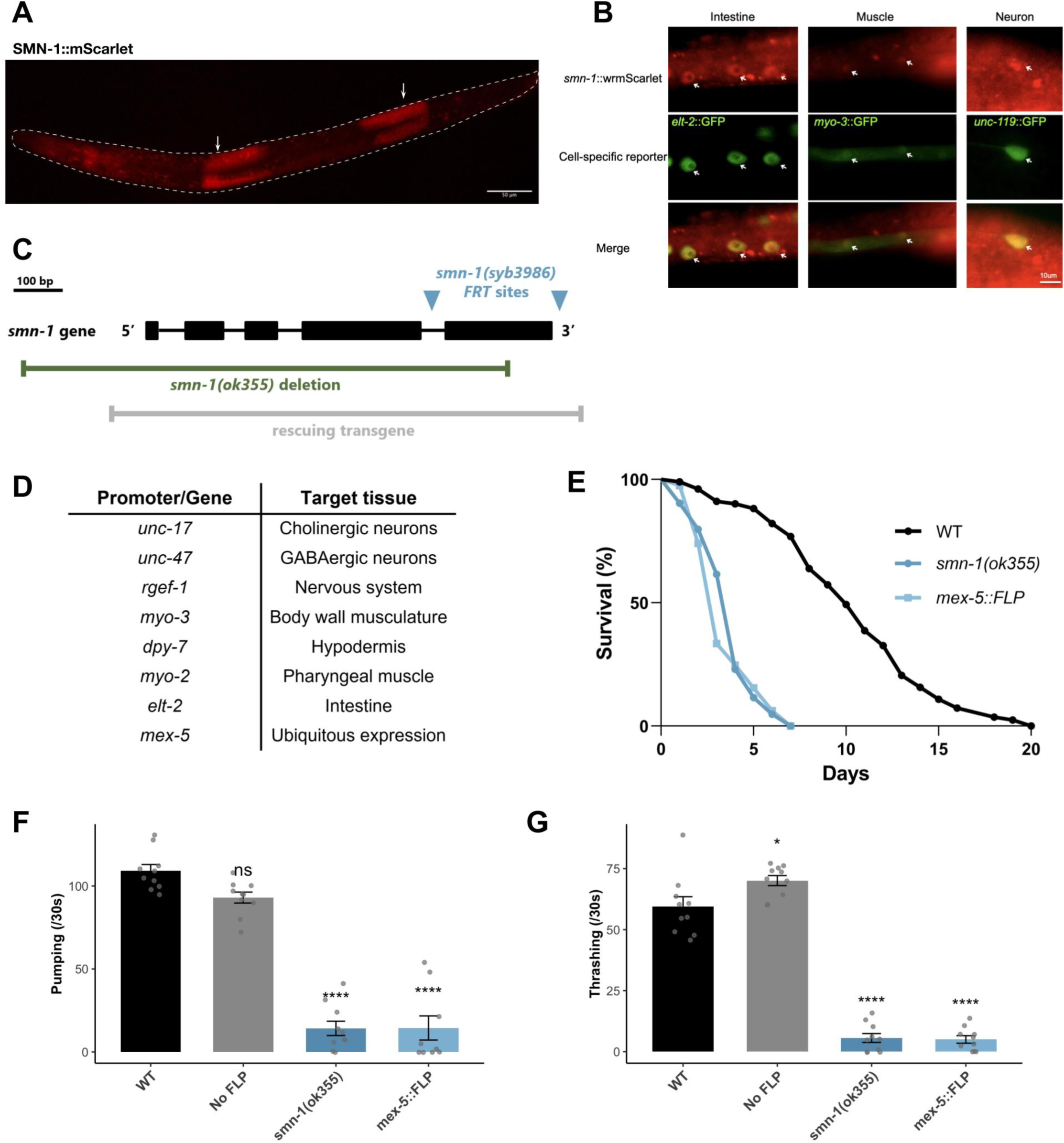
Constructing *smn-1* tissue-specific KOs using the FLP/FRT system. (A) Fluorescence image of animal expressing endogenously tagged SMN-::mScarlet. Dashed outlines indicate animal boundaries, arrows highlight SMN-1 expression in the germline. Scale bar, 50μm. (B) Tissue-specific localization of endogenously tagged SMN-1::mScarlet. SMN-1::mScarlet fluorescence (top) is shown alongside cell-specific GFP reporters (middle) for intestine (*Pelt-2::GFP*), body wall muscle (*Pmyo-3::GFP*), and neurons (*Punc-119::GFP*). Merged images (bottom) demonstrate SMN-1 expression in multiple tissues. Arrows point to representative nuclei or cells. Scale bar, 10 µm. (C) Schematic of the *smn-1* locus showing exon–intron structure, the *smn-1(syb3986)* allele containing FRT sites for tissue-specific excision, the *smn-1(ok355)* null allele, and the rescuing transgene used in this study. (D) Summary of promoters and corresponding target tissues used to drive tissue-specific *FLP* recombinase expression for *smn-1* excision. (E) Kaplan–Meier survival curves of wild-type (n = 100), *smn-1(ok355)* (n = 83), and global KO animals (*mex-5::FLP*) (n = 100) following the 34h post-hatching mark. Global *smn-1* KO (p < 0.0001) using the *FLP*/FRT system phenocopies the shortened lifespan of null *smn-1(ok355)* (p < 0.0001) mutants. Each data point represents the fraction of synchronized animals alive on a given day. (F) Pharyngeal pumping rates (pumps per 30 seconds) measured in wild-type, *smn-1-FRT* control (no *FLP*), *smn-1(ok355)*, and global knockout (*smn-1-FRT; mex-5::FLP*) animals (72h post-hatching). Global *smn-1* KO results in a similar reduction in pumping to *smn-1(ok355)*. (G) Thrashing rates at 120 hours post-hatching. Statistical notes: Individual data points represent single animals; bars show mean ± SEM (n = 10 animals per genotype). Statistical analysis was performed using one-way ANOVA with Dunnett’s multiple comparisons test.

To dissect the tissue- and cell-specific requirements for *smn-1 in vivo*, we used the FLP/FRT system^25,26^ to generate tissue-specific KOs. This system has been demonstrated to achieve high efficiency and specificity, and offers the opportunity to test the requirement for *smn-1* both in specific tissues and in specific neuron types (*e.g.* FLP drivers specific to cholinergic, or to GABAergic, neurons)^26^. Using CRISPR/Cas9 genome editing, we inserted FRT sites flanking exon 5 of the *smn-1* gene (Figure 1C), and crossed this strain with lines expressing FLP recombinase under the control of tissue- and cell-specific promoters (Figure 1D). After constructing these strains, we confirmed FLP-mediated excision of *smn-1* in each line via PCR (Figure S1A).

As an initial test of the efficiency of FLP-mediated *smn-1* deletion *in vivo*, we compared phenotypes of the *mex-5::FLP* line (hereafter “global KO”) to those of the well-characterized *smn-1(ok355)* null allele^18^ (Figure 1C). The *mex-5* promoter drives FLP expression in the germline, and has been shown to induce recombination in 100% of F1 (and subsequent) progeny^25^. We therefore reasoned that these global KO animals should have the same phenotype as *smn-1(ok355)* null mutants if recombination is efficient at the endogenous *smn-1* locus.

Indeed, the global KOs have very similar phenotypes to the null mutants. In both mutants, animals fail to develop past the third larval stage (L3) and therefore the strains must be maintained as balanced heterozygotes. Both null and global KO homozygous mutants also exhibit severely shortened lifespan (Figure 1E). In contrast, animals with *smn-1-FRT* but with no FLP driver have normal lifespans (Figure 1E, S1B). Similar trends are observed for other characteristic *smn-1* phenotypes, including pharyngeal pumping (Figure 1F) and locomotion (Figure 1G). In each case, null and global KOs have severe phenotypic deficits, while *smn-1-FRT* animals without a FLP driver do not. These results confirm that the FLP/FRT system enables efficient KO of *smn-1* and that ubiquitous excision via *mex-5::FLP* resembles null mutants. This system thus provides a robust platform for dissecting the cell-specific roles of SMN-1 *in vivo*.

### *smn-1* is required primarily in the intestine

Having validated the efficiency of the FLP system for generating *smn-1* KOs, we next investigated the tissue- and cell-specific requirements for *smn-1*. We tested KOs in major tissue types—neurons, muscle, intestine, and hypodermis—as well as in subsets of neurons (excitatory cholinergic and inhibitory GABAergic neurons).

We first measured pharyngeal pumping, one of the earliest and most prominent phenotypes observed in *smn-1* null mutants. This behavior, which depends on neuromuscular function, is often used as a proxy for motor decline in *C. elegans* models of SMA^27,28^. Surprisingly, none of the neuronal or muscle-specific KOs cause substantial pumping defects (Figure 2A). In contrast, intestinal KO causes a dramatic reduction in pharyngeal pumping, similar to both the null mutant and the global KO (Figure 2A, 1F).

**Figure 2:**
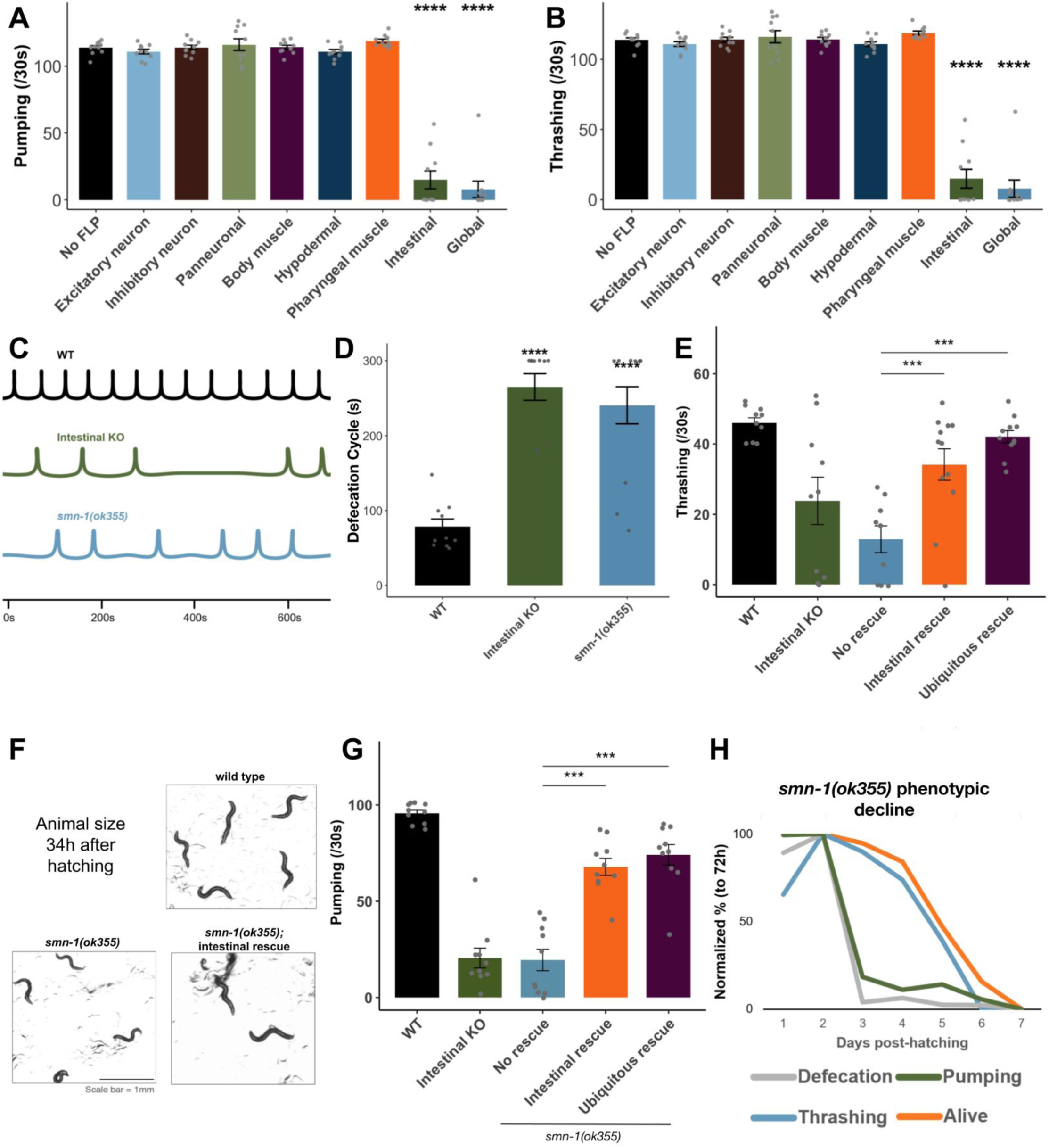
*smn-1* is required primarily in the intestine. (A) Pharyngeal pumping analysis of tissue-specific KOs at 72h post-hatching. (B) Locomotion assay (thrashing) of tissue-specific *smn-1* KOs (120h post-hatching). (C) Defecation cycles for single representative animals at 72h post hatching. (D) Defecation cycle length measured as the interval between posterior body wall muscle contractions (pBoc) in wild-type, intestinal KO and *smn-1(ok355)* animals at 96 hours post-hatching. Intestinal KO and *smn-1(ok355)* mutants exhibit significantly altered cycle periods. (E) Thrashing rates in wild-type, *smn-1(ok355*), and null mutants rescued by intestinal (*Pges-1*), and ubiquitous (*Peft-3*) *smn-1* transgenes. *smn-1(ok355)* mutants exhibit reduced thrashing, which is partially or fully restored upon tissue-specific *smn-1* expression. (F) Representative images of animals at 34h post-hatching. *smn-1(ok355)* mutants display reduced body size, which is partially rescued by intestinal expression. Scale bar, 1 mm. (G) Pharyngeal pumping rates in wild-type, *smn-1(ok355*), and null mutants rescued by intestinal (*Pelt-2*), and ubiquitous (*Peft-3*) *smn-1* transgenes. *smn-1(ok355)* mutants exhibit reduced thrashing, which is partially or fully restored upon tissue-specific *smn-1* expression at 72h post-hatching. (H) Combined analysis of pharyngeal pumping, thrashing, survival, and defecation cycles as a function of time in *smn-1(ok355)* animals. Values are normalized to mutants at 72h post-hatching and are expressed as “normalized percent of phenotype at 72h post-hatching.” Statistical notes: Individual data points represent single animals (or biological replicates as indicated). Bars indicate mean ± SEM. Statistical significance was determined by one-way ANOVA with Dunnett’s multiple comparisons test. Asterisks denote significance (*p < 0.05; **p < 0.01; ***p < 0.001; ****p < 0.0001); ns, not significant.

A second notable defect in *smn-1* null mutants is a decline in locomotory activity. Similar to the pumping phenotype, neither neuronal nor muscle KOs result in locomotory impairment (Figure 2B). However, intestinal KOs again exhibit strong locomotion defects, similar to the null and global KO animals (Figure 2B, 1G). A similar pattern holds for lifespan, in which intestinal KOs, but not other tissue-specific KOs, have substantially shortened lifespan (Figure S1B). Taken together, these findings indicate that phenotypes previously attributed to neuromuscular loss of *smn-1* – progressive defects in locomotion, pumping, and lifespan – are in fact driven by *smn-1* deficiency in the intestine.

We therefore considered whether *smn-1* mutants display additional intestinal defects that might have previously gone undetected. We examined the rhythmic cycle of defecation, which is a behavior responsible for eliminating waste from the lumen of the intestine^30^. Wild-type animals display a stereotyped defecation cycle of approximately once per minute, but *smn-1* null mutants and intestinal KOs exhibit strongly defective defecation behavior, with delayed and irregular cycles (Figure 2C-D).

The conditional KOs indicate that *smn-1* is necessary in the intestine. To test whether intestinal expression of *smn-1* is also sufficient to rescue smn-1 phenotypes, we generated overexpression transgenic lines in which full-length genomic *smn-1* (Figure 1C) is driven by tissue-specific promoters. Intestinal expression rescues mutant phenotypes in both the intestinal KO (Figure S1C and S1D) and in the null mutant (Figure 2E-G). This was further confirmed by the use of two different intestinal promoters (*Pges-1* and *Pelt-2*), both of which rescue null mutant phenotypes (Figure 2E-G, S1D).

Finally, we investigated the temporal dynamics of the mutant phenotypes, which are known to progressively decline^17,18^. We found that both defecation and pumping defects emerge relatively early, whereas locomotive decline occurs later, close to the time of death (Figure 2H). These findings suggest that the initial functional deficits in *smn-1* mutants originate within the alimentary system (intestine and pharynx), and that systemic deterioration — including neuromuscular decline — is a downstream consequence. Together these results indicate that in *C. elegans*, the intestine is selectively vulnerable to loss of *smn-1*. This is in contrast to findings in mammals, which were also assumed to apply to *C. elegans*, where motor neurons are selectively vulnerable to loss of *smn-1*^3,16,17,31,32^.

### Loss of *smn-1* causes specific global splicing defects

Having established that the intestine is the major site of *smn-1* requirement in *C. elegans*, we next explored the molecular consequences of tissue-specific loss of *smn-1*. Given SMN’s essential role in spliceosome assembly and snRNP biogenesis^2,33,34^, we first investigated how deleting *smn-1* in a tissue-specific manner affects splicing. We performed whole-animal RNA-Seq on animals 34 hours post-hatching. We selected this time point as it precedes the onset of overt phenotypic decline, allowing us the possibility of detecting primary transcriptomic changes rather than secondary consequences of systemic dysfunction.

We observed substantial splicing disruption in both global and intestinal *smn-1* KOs. The global KO exhibits a larger number of splicing alterations (Figure 3A), but the overall pattern of splicing defects is similar between the two conditions. The global KO is comparatively enriched for alternative 3′ splice site changes, while the intestinal KO is comparatively enriched for cassette exon skipping.

**Figure 3:**
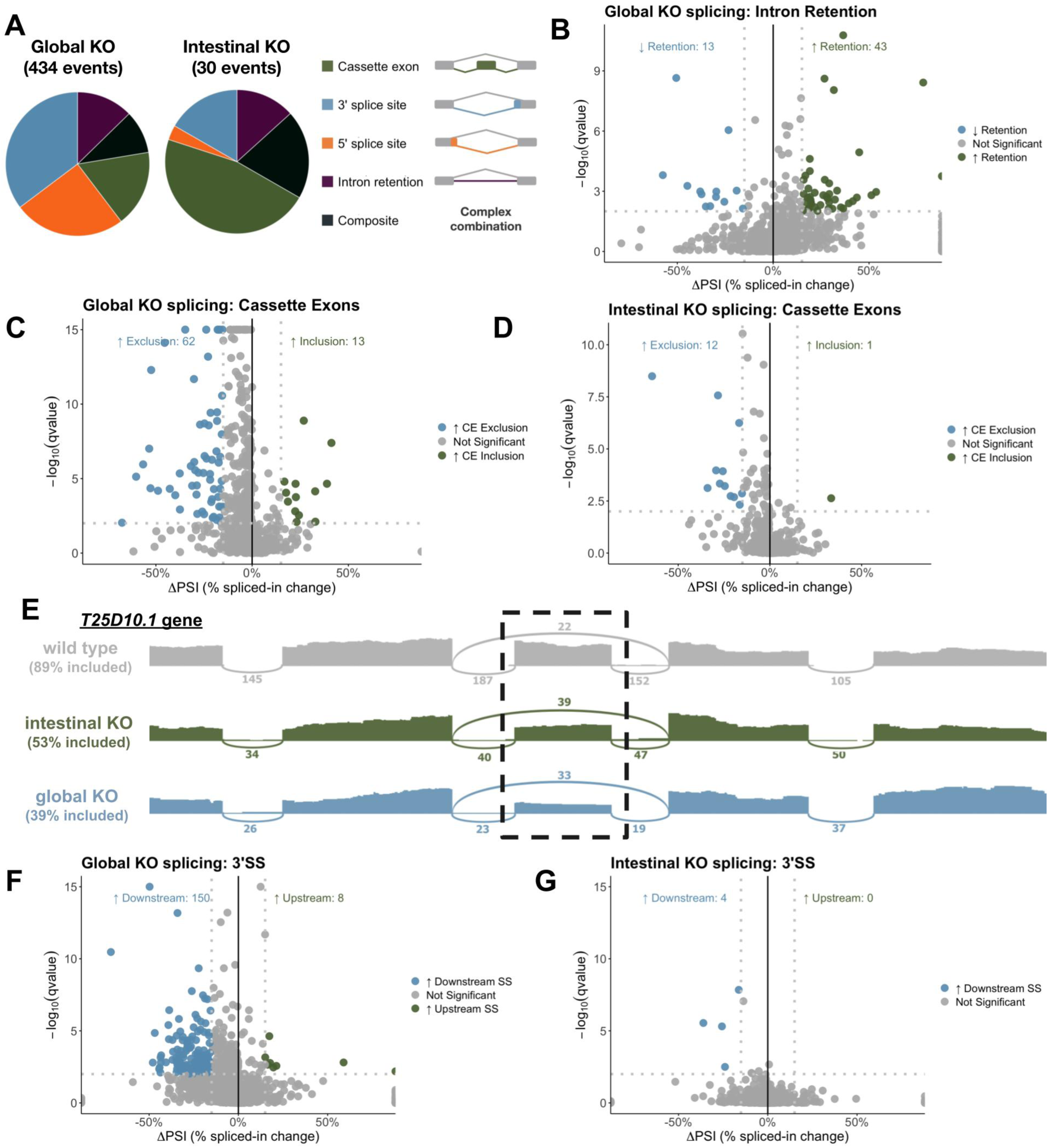
Loss of *smn-1* causes global defects in intron retention, exon inclusion, and 3’ splice site selection. (A) Distribution of dysregulated alternative splicing event types identified by RNA-Seq in global and intestinal KOs. Pie charts show relative proportions of dysregulated splicing types. Total numbers of significant splicing events detected in each condition indicated (ΔPSI > 15% and q ≤ 0.01). Global KO; cassette exons 17.3%, 3’SS 35.3%, 5’SS 25.1%, IR 12.7%, and composite 9.7%. Intestinal KO; cassette exons 46.7%, 3’SS 16.7%, 5’SS 3.3%, IR 13.3%, and composite 20%. (B-D) Volcano plots showing differential splicing events identified by RNA-Seq. (B) and (C) show intron retention and cassette exon events in global KOs, and (D) shows cassette exon events in intestinal KOs. X-axis indicates the change in percent spliced in (ΔPSI), and y axis shows the p-adjusted qvalue (−log10). Dashed lines denote ΔPSI and significance cutoffs (qvalue < 0.01, ΔPSI > 15%). Events with increased intron retention are shown in green (up), decreased intron retention shown in blue (down), and non-significant (p adjusted > 0.01 and PSI < 15%) in gray. Number of significantly increased and decreased intron retention events indicated. (E) Sashimi plot visualization of a representative cassette exon splicing event in the *T25D10.1* gene. Read coverage, splice junction usage, and calculated PSI values indicated for each condition. Dotted rectangle denotes alternative exon. (F, G) Volcano plot of alternative 3′ splice site (3′SS) usage in global KOs (F), and in intestinal KOs. (G) Events with increased usage of downstream or upstream 3′ splice highlighted, using same cutoffs as in panels B-D.

Defective spliceosome assembly might be expected to result in global signatures of splicing failure, such as increased intron retention or increased exon skipping. Indeed, we observe a modest global increase in intron retention (Figure 3B, S2A) and a stronger, widespread increase in exon skipping in both the global and intestinal KOs (Figure 3C-D). In both KOs, exons with increased skipping far outnumber those with increased inclusion (Figure 3C-D). Volcano plots in Figure 3C-D show a leftward tilt (ΔPSI < 0), reflecting a global trend toward exon skipping. This leftward tilt includes many exons with small decreases (gray) and a few exons with pronounced decreases (blue). An illustrative example is shown for the gene *T25D10.1*, in which the central alternative exon is preferentially skipped in both KOs (Figure 3E). Meanwhile the flanking constitutively-spliced exons are spliced normally, suggesting *smn-1* might be particularly important for the splicing of exons with weak recognition elements.

An additional global pattern we observed in *smn-1* mutants is a change in 3’ splice site selection, with increasing usage of downstream 3’ splice sites in *smn-1* mutants (Figure 3F). This pattern is particularly strong for global KOs, but also detectable in intestinal KOs (Figure S2A). No such global trend is apparent for 5’ splice site selection in either mutant (Figure S2A-C). These results indicate that *smn-1* mutation causes large-scale but specific splicing changes, with increased exon skipping and downstream 3’ splice site usage being the most prominent. These observations dovetail with recent findings in both *C. elegans* and in SMA patient-derived cells, where exon skipping and downstream 3’ splice site selection were likewise noted ^35,36^.

We next tested whether neuronal and muscle KOs have similar splicing patterns to those of the intestinal KOs. Indeed, all three tissue-specific KOs show similar global trends. For cassette exons, both muscle and intestinal KOs show the same leftward tilt, indicating a global increase in exon skipping (Figure S2D-F). Likewise for 3’ splice site selection, both KOs have similar trends toward downstream splice site usage (Figure S2G-H). Taken together, these experiments reveal that *smn-1* mutants cause widespread but specific changes in splicing, and that *smn-1* is required for splicing fidelity in multiple tissues. This suggests that the intestine-specific vulnerability to *smn-1* loss is not caused by intestinal specificity of dysregulated splicing, but perhaps rather by tissue-specific downstream responses to loss of *smn-1* and dysregulated splicing.

### Loss of *smn-1* causes up-regulation of a suite of pathogen response and proteostasis genes

We next analyzed gene expression changes that occur in response to loss of *smn-1*, and identified substantial overlap between the global and intestinal KO transcriptomes (Figure 4A), with a high degree of quantitative agreement (Figure 4B), although the global KO causes a substantially broader set of transcriptomic changes (Figure 4A).

**Figure 4:**
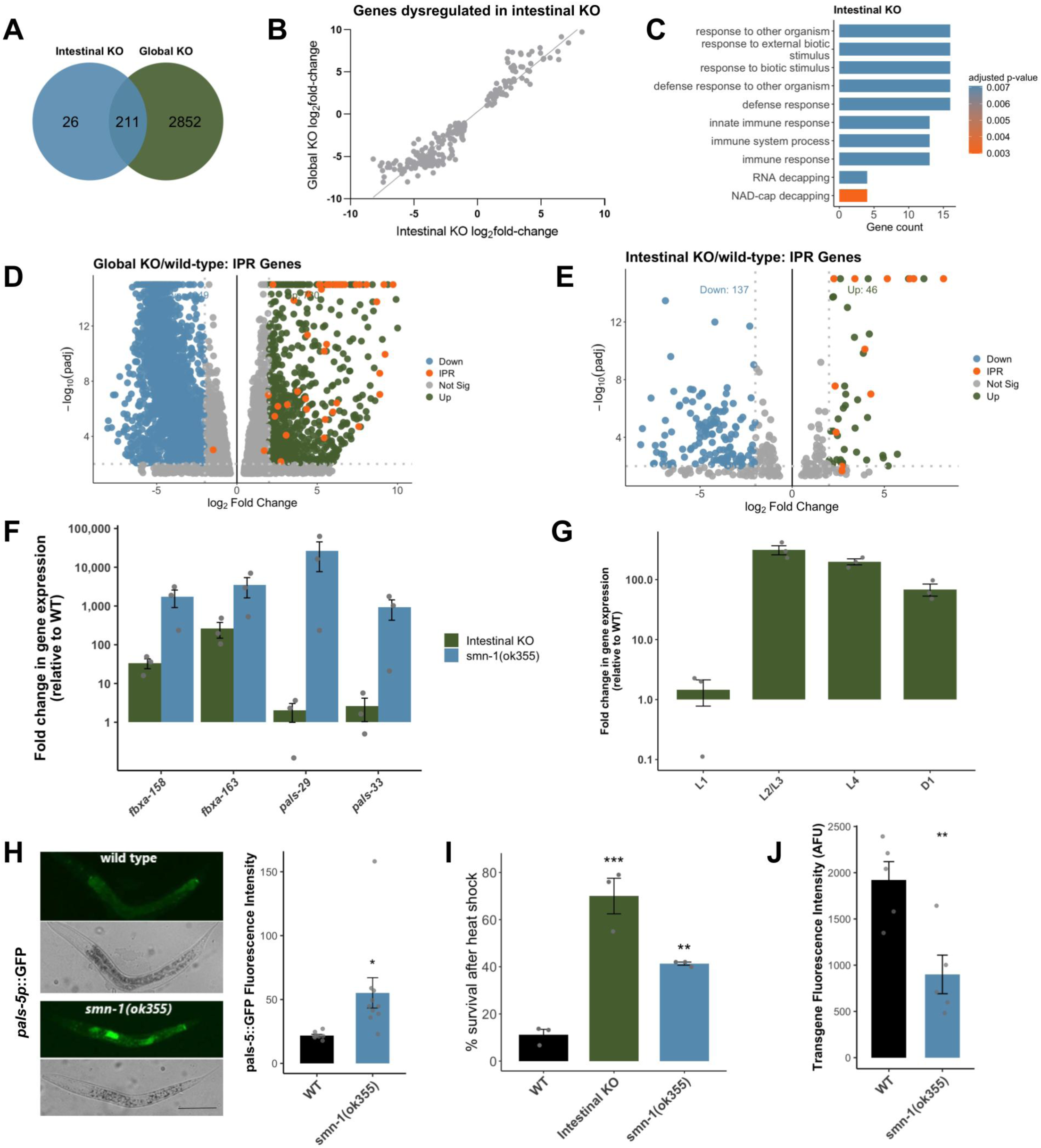
Loss of *smn-1* causes up-regulation of a suite of pathogen response and proteostasis genes. (A) Venn diagram showing the overlap of differentially expressed genes (*p* <0.05 and log2fold change > 1.5) identified by RNA-Seq in intestinal and global KOs. (B) Scatter plot including all genes undergoing differential expression in intestinal KOs. The plot compares log2 fold changes in the intestinal KO versus those in global KO. Each point represents an individual gene, illustrating concordance of transcriptional responses between tissue-specific and global SMN loss. Pearson’s correlation coefficient (R^2^) value = 0.911. C. Gene Ontology (GO) enrichment analysis of genes dysregulated in intestinal KO. Enriched biological process terms are shown, with bar length indicating gene count and color representing adjusted p-value. (D, E) Volcano plots of differentially expressed genes in global (D) and intestinal KO (E). Log2 fold change is plotted against −log10 adjusted p-value. Genes associated with the Intracellular Pathogen Response (IPR) are highlighted (orange), with significantly upregulated (green) and downregulated (blue) genes indicated. (F) qPCR for selected IPR and stress-responsive genes in intestinal KOs and *smn-1(ok355)* mutants relative to wild type. (G) qPCR of *fbxa-163* gene expression throughout various larval developmental stages of intestinal KO mutants. Upregulation of *fbxa-163* precedes the appearance of apparent mutant phenotypes. (H) (Left) Representative fluorescence and corresponding brightfield images of *pals-5p::GFP* transcriptional reporter expression in wild-type and *smn-1(ok355)* animals two days post-hatching, showing elevated reporter activity upon *smn-1* loss. (Right) Quantification of *pals-5p::GFP* fluorescence intensity. Each dot represents an individual animal; bars indicate mean ± SEM. Scale bar = 100 μm. (I) Survival of wild-type, intestinal KO, and *smn-1(ok355)* animals following acute heat shock at 37.5°C followed by overnight recovery at 20°C (34h post-hatching). Data demonstrates enhanced heat stress resistance in intestinal KOs and null mutants. (J) Quantification of transgene fluorescence intensity in wild-type and *smn-1(ok355)* animals two days post-hatching, showing increased transgene *(otIs381[ric-19prom6::NLS::gfp]*) silencing in null mutants. Statistical notes: Individual data points represent single animals (or biological replicates as indicated). Bars indicate mean ± SEM. Statistical significance was determined by one-way ANOVA with Tukey’s or Dunnett’s multiple comparisons test, or by Student’s t-test where indicated. Asterisks denote significance (*p < 0.05; **p < 0.01; ***p < 0.001; ****p < 0.0001); ns, not significant.

Upregulated genes in both mutants are enriched for immune-related gene ontology categories (Figure 4C, S3A). Indeed, we found that many of the top upregulated genes fall into a single, well-characterized pathway: the Intracellular Pathogen Response (IPR)^37^. This response, triggered by natural intracellular infections of the intestine, induces a large suite of genes involved in proteostasis and immunity—particularly E3 ubiquitin ligases such as the *fbxa* gene family^37–39^. Many of these upregulated IPR genes are expressed exclusively in the intestine^37–39^. Labeling IPR genes on the gene expression volcano plots (Figure 4D-E, orange) reveals global upregulation, with many genes being upregulated 100-fold or more (Figure 4D-E).

In contrast, analysis of muscle and neuronal *smn-1* KOs reveals very few changes in gene expression (only 1 gene upregulated in the neuronal KO, 0 in the muscle KO) (Figure S3C-D). Thus, although splicing dysregulation is similar in all three tissue-specific KOs (Figure S2A-G), only the intestinal KO responds with large-scale transcriptional responses (Figure 4E). We suspect that this is because the IPR genes are poised to be activated only in the intestine, and not in muscles or neurons.

qRT-PCR analysis confirms a dramatic induction of IPR genes in *smn-1* mutants (Figure 4F), with some E3 ubiquitin ligase genes (e.g., *fbxa-158*, *fbxa-163*) upregulated by hundreds- to thousands-fold in both KOs. Notably, while downstream effectors of the IPR (such as *fbxa* genes) are strongly upregulated in both KOs, upstream regulators of the pathway (e.g., *pals* genes, whose biochemical function is currently unknown) are induced in the global KO but not the intestinal KO (Figure 4F).

We next assessed the temporal dynamics of IPR activation. qRT-PCR across larval development reveal that *fbxa-158* and *fbxa-163* are highly expressed beginning in early larval development (Figure 4G, S3B), well before the onset of phenotypic decline in *smn-1* mutants. This suggests that these IPR gene expression increases are not merely the consequence of a general decline of health, but might constitute upstream contributors to *smn-1* mutant phenotypes.

Given the robust activation of IPR genes in the *smn-1* KOs, we next asked whether *smn-1* mutants exhibit hallmarks of IPR-associated phenotypes. A transcriptional *pals-5::GFP* transgene is often used as a readout for IPR activity, with increased intestinal GFP observed upon infection or genetic activation of IPR^37,39,40^. We find that *smn-1* mutants also result in robust activation of *pals-5::GFP,* and that this activation is confined to the intestine (Figure 4H).

IPR activation causes resistance to stressors (*e.g.* heat stress, infection) but at the expense of shortened lifespan and developmental delay^41^. We already observed that *smn-1* mutants have shortened lifespan and development delays (Figure 1), and we next tested whether *smn-1* mutants, like IPR-active mutants, are resistant to heat stress^37,39^. Indeed, both the global and intestinal KOs are significantly resistant to heat stress-induced lethality (Figure 4I). IPR activation also causes increased small RNA biogenesis, a readout of which is increased silencing of repetitive transgenes^42^. *smn-1* mutants, like other IPR-active mutants, have this transgene-silencing phenotype as well (Figure 4J). Taken together, these results demonstrate that *smn-1* mutation leads to strong overexpression of IPR genes accompanied by the hallmark phenotypes of the IPR, including increased stress resistance at the expense of developmental and health defects.

### *smn-1* mutation causes increased proteasome activity

Many of the most highly-upregulated IPR genes are members of the ubiquitin proteostasis pathway. In particular, many E3 ubiquitin ligases (such as members of the *fbxa* family of genes) are highly upregulated upon IPR activation, as well as upon *smn-1* KO. It is thought that these E3 ubiquitin ligases might directly target pathogenic proteins with ubiquitin for destruction by the proteasome as an innate immune response^43–45^. We hypothesized that in *smn-1* mutants, the overexpression of these proteostasis genes drives elevated proteasome activity even in the absence of infection, contributing to tissue dysfunction.

To test this, we used a Ub(G76V)::GFP transgene as an *in vivo* reporter of proteasome activity^37^. In this system, GFP is fused to a modified ubiquitin that resists deubiquitination, thus targeting it for constitutive degradation by the proteasome (Figure 5A). Inhibiting proteasome activity with the drug Bortezomib (BTZ) at 10 μM prevents degradation of this transgenic protein, leading to strong GFP fluorescence (Figure 5A-B). However, in *smn-1* mutants, this BTZ treatment fails to increase GFP levels (Figure 5A-B). This is consistent with our hypothesis that *smn-1* mutants possess elevated basal proteasome activity, such that even in the presence of a proteasome inhibitor, ubiquitinated GFP is effectively degraded. At higher BTZ concentration (50 μM), however, both wild-type and *smn-1* mutants exhibit elevated GFP accumulation (Figure 5A), suggesting that elevated BTZ concentration is sufficient to overcome high basal levels of proteasome activity in *smn-1* mutants.

**Figure 5:**
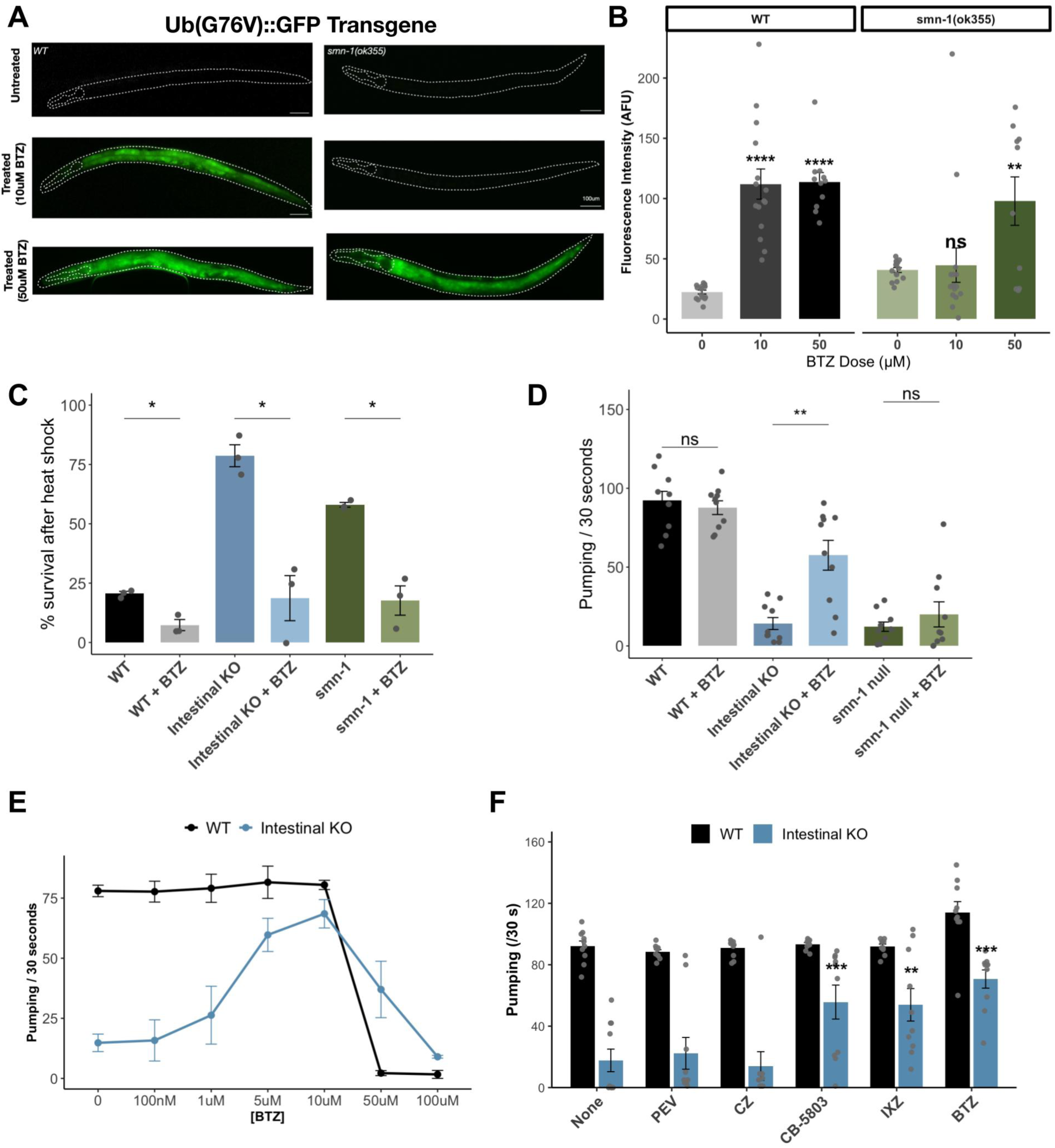
*smn-1* mutation causes increased proteasome activity which contributes to the *smn-1* mutant phenotypes. (A) Representative fluorescence images of animals expressing an i*n vivo* transgene to visualize UPS activity. Images of *smn-1(ok355)* and wild-type worms (34h post-hatching) expressing *Ub(G76V)::GFP* from an integrated transgene (*mgIs77[rpl-28p::ub(G67V)::GFP*) showing treatment with and without treatment with the proteasome inhibitor bortezomib (BTZ) at two concentrations (10uM and 50uM) for 8 hours. Dashed outlines indicate animal boundaries. Scale bar, 100 µm. (B) Quantification of *Ub(G76V)::GFP* fluorescence intensity shown in A (n = 15 per genotype) under untreated conditions or following bortezomib treatment (BTZ; 0μM, 10μM, 50μM). (C) Survival following acute heat shock in the presence or absence of BTZ (10μM). Bars demonstrate differential effects of proteasome inhibition on thermotolerance across genotypes (34h post-hatching). Survival data were analyzed by pairwise comparisons between untreated and BTZ-treated animals within each genotype using two-sided Student’s t-tests. (D) Pharyngeal pumping rates measured with or without 10μM BTZ treatment. Partial rescue of pumping behavior in intestinal KO animals upon proteasome inhibition. Pumping rates were compared between untreated and BTZ-treated animals within each genotype using two-tailed unpaired Student’s t-tests. (E) Left: Dose–response analysis of pharyngeal pumping rates in wild-type and intestinal KOs with increasing concentrations of BTZ at 72h post-hatching. (F) Comparison of pharyngeal pumping rates in wild-type and intestinal KOs following treatment with distinct proteostasis-modulating compounds (pevonedistat, carfilzomib, CB-5083, ixazomib, and bortezomib) at a concentration of 10μM. Statistical notes: Individual data points represent single animals (or biological replicates as indicated). Bars indicate mean ± SEM. Statistical significance was determined by one-way ANOVA with Tukey’s or Dunnett’s multiple comparisons test, or by Student’s t-test where indicated. Asterisks denote significance (*p < 0.05; **p < 0.01; ***p < 0.001; ****p < 0.0001); ns, not significant.

We next asked whether hyperactive proteasome activity might contribute to *smn-1* mutant phenotypes by testing whether pharmacological proteasome inhibition is sufficient to reverse *smn-1* mutant defects. Indeed, BTZ suppresses the elevated heat-shock resistance observed in *smn-1* KOs, restoring survival rates toward wild-type levels (Figure 5C). Likewise, BTZ treatment significantly improves pharyngeal pumping in intestinal *smn-1* KOs (Figure 5D), while in the global KO it causes an increase in pharyngeal pumping that is not statistically significant.

Dose–response assays reveal that pharyngeal pumping phenotypes improve with increasing BTZ concentrations up to 10 μM, at which point intestinal KOs are restored to near wild-type levels (Figure 5E). At 50 μM, BTZ treatment causes wild-type worms to be very sick and exhibit nearly 0 pharyngeal pumping (Figure 5E). In contrast, intestinal KOs treated with 50 μM BTZ not only tolerate it, but have higher pumping rates than untreated animals (Figure 5E).

To further test how elevated proteasome activity contributes to intestinal KO phenotypes, we examined the effects of pharmacological inhibitors targeting distinct steps of the ubiquitin–proteasome system (Figure 5F). In wild-type animals, none of the compounds substantially alter pharyngeal pumping rates. In contrast, intestinal *smn-1* KOs show significant improvement in pumping upon treatment with inhibitors targeting the 20S proteasome core^46,47^, including BTZ and ixazomib (IXZ) (Figure 5F). The protein p97/VCP binds to ubiquitinated substrates and facilitates subsequent proteasomal degradation^48^. Inhibition of p97/VCP with the drug CB-5803 rescues pumping defects as well (Figure 5F). On the other hand, compounds acting upstream in the ubiquitin pathway, including the deubiquitinase inhibitor capzimin (CZ) and the NEDD8-activating enzyme (E3 ubiquitin ligase activator) inhibitor pevonedistat, do not rescue (Figure 5F). These results indicate that excessive proteasome activity contributes to the functional defects caused by intestinal loss of *smn-1*.

### Upregulated *fbxa* genes contribute to the *smn-1* mutant phenotype

We next asked whether we could identify specific elements of the *smn-1* mutant transcriptome responsible for specific elements of the *smn-1* mutant phenotypes. We focused on the *fbxa* genes, due to their high levels of overexpression and their function as E3 ubiquitin ligases. This positions them as potential mediators of the hyperactive proteolysis observed in *smn-1* mutants, due to increased ubiquitin ligation of their target proteins.

In *C. elegans* the *fbxa* gene family is greatly expanded (200+ members), and *fbxa* genes tend to be genomically clustered^42, 45^. We observed that *fbxa* upregulation in *smn-1* mutants is not global across all clusters – some clusters (*e.g.* Chr II: 1.6 Mb) exhibit high overexpression in both the global and intestinal KOs (Figure 6A), while others (*e.g.* Chr III: 0.9 Mb) show no significant change (Figure 6B). This specificity suggests a focused transcriptional program rather than a global derepression of *fbxa* genes.

**Figure 6:**
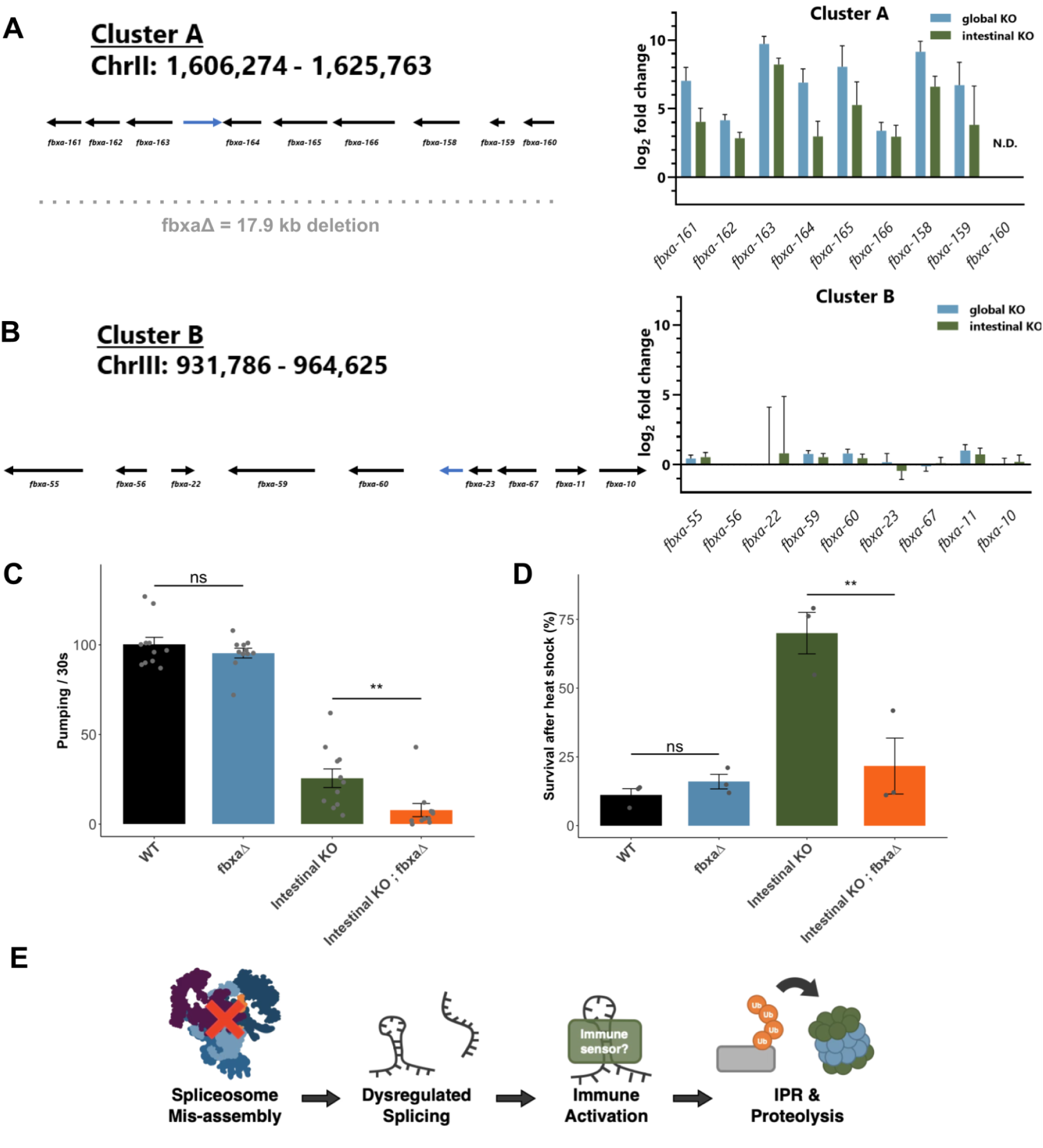
Upregulated *fbxa* genes contribute to the *smn-1* mutant phenotype. (A, B) Genomic organization and expression analysis of *fbxa* (A) gene Cluster A on chromosome II (ChrII: 1,606,274–1,625,763) and (B) gene Cluster B on chromosome III (ChrIII: 931,786–964,625). Arrows indicate gene orientation. RNA-Seq–derived log2 fold changes for individual *fbxa* genes are shown for global and intestinal KOs relative to wild-type. In (A), a CRISPR-mediated 17.9 kb deletion spanning the *fbxa* cluster (*fbxaΔ*) is indicated. Error bars represent SEM; N.D., not detected. (C) Pharyngeal pumping rates showing no rescue of the intestinal KO by the *fbxaΔ* deletion. (D) Survival following acute heat shock, showing that the intestinal KO is rescued by the *fbxaΔ* deletion (34h post-hatching). Statistical significance was determined by unpaired two-tailed t-tests for specific comparisons: *fbxaΔ* vs WT, and *fbxaΔ ; smn-1-FRT ; elt-2::FLP* vs Intestinal KO. (E) Working model illustrating how SMN loss leads to spliceosome mis-assembly and dysregulated splicing, resulting in activation of innate immune signaling pathways and induction of the intracellular pathogen response (IPR) which includes increased ubiquitin-mediated proteolysis. Statistical notes: Individual data points represent single animals (or biological replicates as indicated). Bars indicate mean ± SEM. Statistical significance was determined by one-way ANOVA with Tukey’s or Dunnett’s multiple comparisons test, or by Student’s t-test where indicated. Asterisks denote significance (*p < 0.05; **p < 0.01; ***p < 0.001; ****p < 0.0001); ns, not significant.

To test whether *fbxa* upregulation contributes to the *smn-1* mutant phenotypes, we used CRISPR/Cas9 to delete all ten genes in the *fbxa* cluster at Chr II:1.6 Mb (Figure 6A). This is the most highly-overexpressed *fbxa* cluster, the genes of which are known to be expressed specifically in the intestine^49,50^. We crossed this *fbxaΔ* mutant into the intestinal KO background, and assessed pharyngeal pumping (Figure 6C), finding that the *fbxa* deletion does not rescue *smn-1*.

On the other hand, *fbxa* deletion does significantly suppress the enhanced heat stress resistance phenotype of the *smn-1* intestinal KO, reducing it to near wild-type levels (Figure 6D). These results are similar to previous experiments in which IPR-activating genes were found to promote heat stress resistance mediated by upregulation of specific *fbxa* genes^38^. Together our results indicate that specific *smn-1* phenotypes (heat-stress resistance) are driven by specific upregulated components of the proteostasis gene network (*fbxa* genes). Meanwhile other phenotypes (*e.g.* pharyngeal pumping defects) are also mediated by elevated proteasome activity, but are not driven solely by the *fbxa* genes identified here.

## DISCUSSION

### Cell-specific vulnerabilities to loss of *smn-1*

Many ubiquitously expressed genes cause highly selective neurodegenerative diseases, such as Huntington’s disease and Amyotrophic Lateral sclerosis^14,15^. The basis for this cell-specific vulnerability remains unclear. We set out to address this question for *smn-1* in *C. elegans*. We show that *smn-1* is indeed required in a cell-specific manner, but unexpectedly not in neurons, but rather in the intestine. Both survival defects and behavioral phenotypes originate from intestinal loss of *smn-1.* This is striking in the context of mammalian studies, which clearly demonstrate motor neuron–specific vulnerability to *SMN* loss^3,51–53^.

It is also surprising in the context of *C. elegans* studies where motor-neuron vulnerability is assumed to be the case^17,18,54–56^. At least one study has obtained experimental results to this effect, where neuronally-expressed *smn-1* partially rescued a null mutant^18^. One potential explanation for this is the differing nature of the rescuing transgenes used. We used the endogenous *smn-1* 3 ’UTR, while it appears the previous study used a heterologous 3 ’UTR. The most common such 3 ’UTR is taken from *unc-54*, and is known to permit ectopic intestinal expression^57^. It is therefore possible that the reported rescue reflected both intestinal and neuronal expression, in which case the results would be consistent with those presented here. In any event, we show here that *smn-1* intestinal KO causes defects similar to null mutants, while intestinal expression, tested with two different promoters, rescues null mutant defects. Worm models of SMA therefore differ from mammalian models with respect to cell-specific vulnerability.

Our question then changes to: why is the intestine selectively vulnerable to loss of *smn-1* in worms? One potential answer is the unique role the intestine plays in sensing intracellular pathogens. As the first line of defense against such pathogens, many IPR genes are expressed only in the intestine. Therefore, in *smn-1* null mutants, even though all cells experience aberrant spliceosome activity, only the intestine responds to such RNA dysregulation by deploying IPR gene expression. In the absence of actual infection, this IPR response is maladaptive, causing aberrantly high proteasome activity and resulting in shortened lifespan, decreased pharyngeal pumping, and increased heat-stress resistance.

What might these results tell us about human SMA? Two potential lessons may apply. The first is the importance of SMN in non-neuronal cells. Although motor neurons are the primary disease-relevant vulnerability in SMA, other tissues exhibit defects as well^10,12,58^. It may be that as patients survive longer due to new motor neuron-preserving therapies, these defects will become increasingly important. The second potential lesson is that the cell-specific proteostasis environment is an important determinant of cell-specific vulnerabilities. Recent work in zebrafish showed that *in vivo* rates of proteasome-mediated degradation are high in spinal motor neurons, and are further influenced by TDP-43, a driver of motor-neuron specific degeneration in ALS^59^. Human SMA neurons also exhibit particularly high levels of unfolded protein response genes^60^. Taken together, these observations suggest that cell-specific baseline proteostasis states may predispose certain tissues—worm intestine or vertebrate motor neurons—to degeneration.

### Proteasome inhibition improves *SMN* phenotypes

Why might defects in spliceosome assembly lead to upregulation of IPR proteostasis genes? We speculate that global splicing defects (Figure 3) generate aberrant RNA products that are detected by the cell as evidence of intracellular infection. Indeed, IPR activation by Orsay virus infection is thought to depend on sensing RNA viral replication intermediates such as dsRNA^61^. If accumulation of mis-spliced endogenous RNA is sensed by the same machinery, this could drive the maladaptive activation of IPR and proteostasis genes observed here (Figure 6E).

Although the IPR appears to be a nematode-specific immune pathway^62^, similar immune-response mechanisms may operate in other SMA models. Recent work in *Drosophila* has demonstrated that innate immunity genes, including several ubiquitin-proteasome genes, are upregulated in *Smn* mutants^63^.

Comparable patterns do not appear to have been described in mammalian studies, although various immune defects have been identified^64^.

Proteasome inhibition may therefore represent an effective strategy for ameliorating phenotypes caused by maladaptive increases in proteostasis gene expression in SMN mutants, and perhaps in other disorders of RNA dysregulation. BTZ administration markedly improves health in our *C. elegans* mutants, and there is some evidence for similar effects in mammals: in a mouse SMA model, BTZ treatment improved select phenotypes, including motor function^65^. This effect was originally interpreted as resulting from increased SMN protein levels, but a plausible alternative explanation is rescue via global proteasome inhibition. It will be interesting in the future to test the extent to which our observations are broadly applicable: first, do other states of dysregulated RNA (*e.g.* mutations in splicing factors or other RNA binding proteins) elicit similar responses? And second, are similar responses elicited by RNA dysregulation in other organisms, including in human RNA-mediated pathologies?

## DATA AVAILABILITY

Raw RNA-Seq reads have been deposited at NCBI GEO, under accession number GSE278038.

## Materials and Methods

### C. elegans strains and maintenance

*C. elegans* strains were maintained on nematode growth medium (NGM) plates seeded with *E. coli* OP50 at 20 °C under standard conditions unless otherwise noted, as previously described by Brenner (1974)^66^. Wild-type animals correspond to the Bristol N2 strain. A complete list of strains used in this study: N2, *smn-1(ok355) I/hT2[bli-4(e937) let-?(q782) qIs48] (I;III)*, *smn-1-FRT*, *smn-1-FRT;elt-2::FLP*, *smn-1-FRT;mex-5::FLP, smn-1-FRT;rgef-1::FLP, smn-1-FRT;myo-3::FLP, smn-1-FRT;unc-119::FLP, smn-1-FRT;unc-47::FLP, smn-1-FRT;dpy-7 ::FLP, smn-1(ok355) ; Ex[Pelt-2::smn-1], rpl-28p::ub(G76V)::gfp, otIs381[ric-19prom6::NLS::gfp]*.

The *smn-1(ok355)* deletion allele was generated by the *C. elegans* gene knockout consortium^67^ and used as a loss-of-function model of spinal muscular atrophy, consistent with prior studies in *C. elegans*^18,68^ . Developmental staging was performed by synchronization via hypochlorite treatment, and animals were analyzed at the indicated larval or adult stages.

### CRISPR/Cas9-mediated generation of FRT sites

*smn-1-FRT* and *fbxaΔ* mutant strains were generated by SunyBiotech. FRT sites were introduced into the endogenous *smn-1* locus flanking exon 5 using CRISPR/Cas9-mediated genome editing, and correct insertion of FRT sites was confirmed by Sanger sequencing. The resulting allele is referred to as *smn-1(syb3986)*. A multi-gene deletion spanning approximately 17.9 kb of the *fbxa* gene cluster on chromosome II was also generated using CRISPR/Cas9, and deletes the genes *fbxa-161, fbxa-162, fbxa-163, btb-15, fbxa-164, fbxa-165, fbxa-166, fbxa-158, fbxa-159,* and *fbxa-160*. Precise deletion was validated by PCR using primers outside the targeted region and validated by sequencing, as well as by Sanger sequencing. We refer to this allele as *fbxaΔ*.

### Transgenic lines

Rescue constructs were generated using the entire *smn-1* genomic region, including introns, from start codon to 3’ UTR. Plasmids were injected at 50 ng/μl. The same genomic insert was cloned into multiple promoter drivers (*rgef-1, myo-3*, *ges-1*, *elt-2*, *eft-3*) via Gateway LR recombination.

### Lifespan analysis

Lifespan assays were performed as previously described^29^. Synchronized L4 larvae (34h post-hatching) were transferred to fresh NGM plates seeded with OP50 (no FuDR used) and scored daily for survival. Animals that failed to respond to gentle prodding were scored as dead. To avoid overcrowding due to progeny production and to ensure access to sufficient food, animals were transferred to new plates when considered necessary. Animals that crawled off the NGM plate were excluded from the data.

Survival curves were generated using Prism, and statistical significance was assessed using the log-rank (Mantel–Cox) test.

### Defecation Assay

Observing the tail end of the worm at each developmental stage and timing the seconds between each defecation. The defecation can be observed by the contraction of the tail and excretion. Performed for about a total of 50 worms for each strain, 10-15 worms per experiment. Calculations were converted to ‘defecations per hour’ with the following equation: (60*60)/Defecation time in seconds.

### Pharyngeal pumping assay

Pharyngeal pumping assays were performed at various larval stages. Grinder movement in any axis was scored as a pumping event and measured for 30 seconds. The average pumping rates (±SEM) were combined from at least three independent trials (*n* > 20 animals in total/genotype).

### Motility (thrashing) assay

Animals were raised as in lifespan assays. On specific days, animals were removed from NGM plates, suspended individually in M9 medium, and the number of body bends in a 30s period was counted. One body bend was counted every time the part of the worm just behind the pharynx reached a maximum bend in the opposite direction from the bend last counted. Data was analyzed using Microsoft Excel.

### Thermotolerance

Thermotolerance assay was performed as described previously^38^. Worms were grown on standard NGM plates until the L4 stage at 20°C. Around ∼100 L4 worms were picked onto fresh plates and were shifted to 37.5°C for two hours. Following heat treatment, plates were placed on the bench in a single layer for 30 minutes to recover, and then moved to a 20°C incubator overnight. Worms were scored for survival 24 hours after heat shock. Animals not pumping or responding to touch were scored as dead. For each strain, three independent assays were performed in triplicate, with at least 20 worms per plate

### Drug treatments

Bortezomib (Sigma Aldrich) was dissolved in dimethylsulfoxide (Sigma Aldrich) and then added to molten NGM agar to the desired concentration before pouring into petri dishes. Control NGM plates contained the respective dilution of the vehicle. Plates were allowed to dry for 1 day before seeding with OP50 bacteria, after which they were incubated at room temperature. Staged worms were then transferred onto the drug/control plates and scored as per the behavioral assay. Each condition was run in triplicate (n > 20 animals in total/genotype).

### Intestinal barrier assay (Smurf)

Animals were raised as described above for lifespan assays. Intestinal permeability assay was performed as previously described ^29,69^.

### Microscopy Analysis

Fluorescence imaging was performed using a Zeiss AxioImager Z1 equipped with 20X air objective and, in some instances, a 63X oil immersion objective. Animals were immobilized using 50mM sodium azide and mounted on 2% agarose pads for imaging. Image processing and quantification were performed using Fiji/ImageJ. Fluorescence intensity measurements were background-subtracted and normalized as indicated.

### RNA-Seq and data processing

*smn-1(ok355)* homozygous mutants arrest at late larval stages and were maintained as heterozygotes using the balancer hT2(I;III). For this reason, all mutants and *wild-type* animals were synchronized by bleaching and allowed to grow for 2 days (around 36 hours) post-hatching for sequencing purposes (larval stage 4 for wild-type animals). For *smn-1(ok355)* and *smn-1-FRT* ; *mex-5 :: FLP* homozygotes, ∼2500 individuals were picked for each biological replicate under a fluorescence dissecting microscope. Stage matched mid-L3 stage animals that had been raised at 20 °C were used for total RNA isolation with Direct-zol RNA miniprep (Zymo Research).

RNA-Seq was performed using NEBNext® Ultra™ II RNA Library Prep Kit for Illumina. The constructed libraries were sequenced as 150 bp paired-end reads on an Illumina platform. Reads were mapped to the worm transcriptome using STAR^70^, gene expression was analyzed using DESeq2^71^ and alternative splicing analyzed by JUM^72^, using standard parameters as previously described.

### GO term enrichment analysis

Gene Ontology (GO) enrichment analysis was performed using the clusterProfiler package in R. Differentially expressed genes were analyzed for enrichment of Biological Process (BP) terms using the *enrichGO* function with *C. elegans* annotations from org.Ce.eg.db. Gene symbols were used as identifiers, and p-values were adjusted for multiple testing using the Benjamini–Hochberg method. GO terms with an adjusted p-value < 0.05 were considered significantly enriched.

### Quantitative RT-PCR

cDNA was synthesized 500ng RNA using Verso cDNA Synthesis Kit (ThermoFIsher). qRT-PCR was performed using SYBR chemistry using PowerUp SYBR Green Master Mix on a BioRad thermocycler. Relative expression was calculated using the ΔΔCt method, normalized to actin. Primers used were as follows: *fbxa-158*F 5’ GCGCCGATCCAGTTCCAAAG 3’, *fbxa-158*R 5’ GAACAACGCTCCGTAAGTTTCG 3’, *fbxa-163*F 5’ TCGAAGCGTCAATCCAGTTCC 3’, *fbxa-163*R 5’ ACGACAAGACTTTCGATCACAC 3’, *pals-29*F 5’ CAAAGATGGCTTCAGGCGCAG 3’, *pals-29*R 5’ GCATGTCTCTTGGCTATTTCTTGG 3’, *pals-33*F 5’ GAGTCACAGATTCAACTTCACG 3’, *pals-33*R 5’ CTGATTGTTGATTGTGCTCGTAGG 3’.

## SUPPLEMENTAL FIGURES

**Figure S1:**
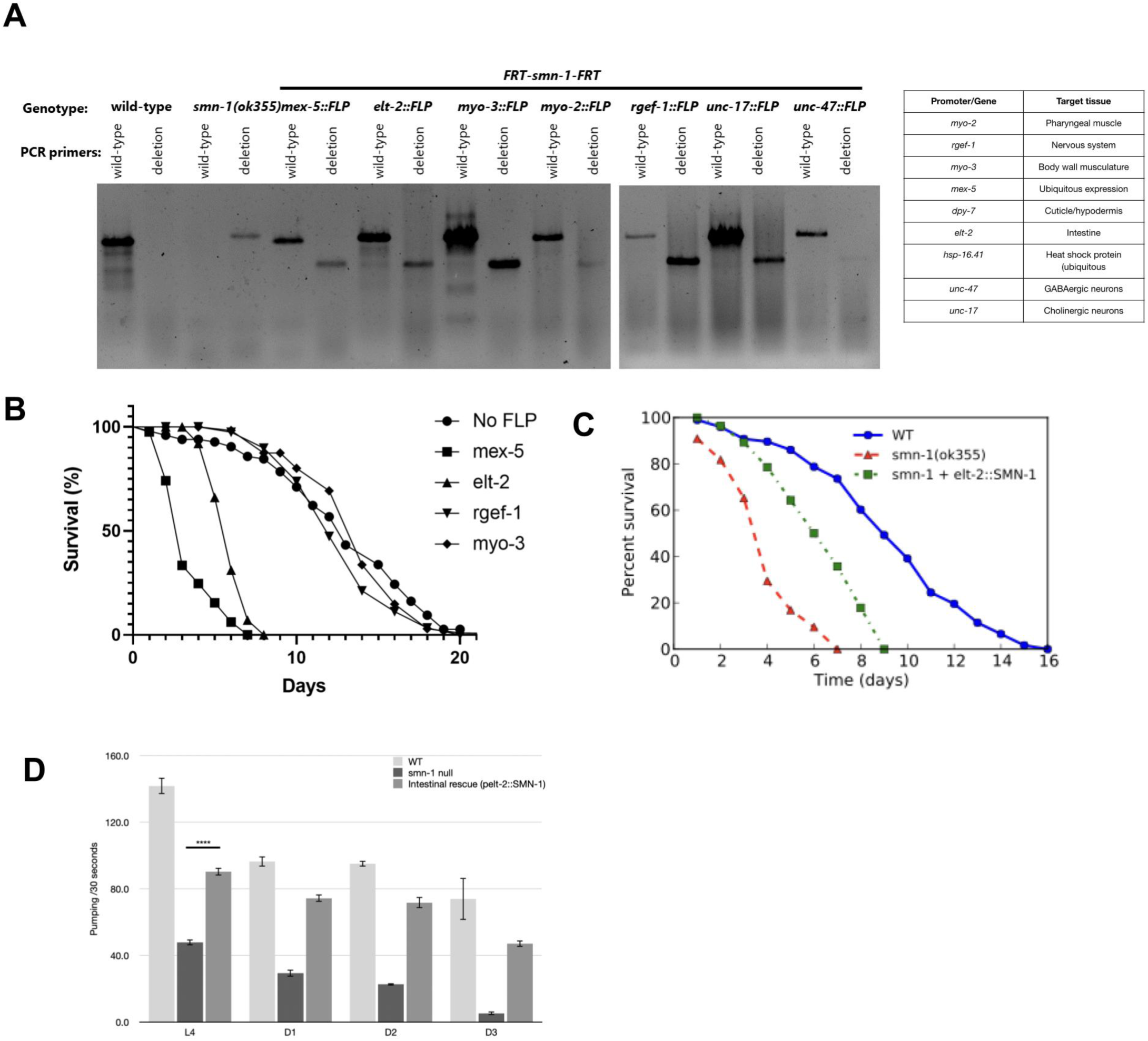
Validation of tissue-specific smn-1 excision and additional phenotypic analyses. (A) PCR-based validation of tissue-specific excision of the *smn-1-FRT* allele in animals expressing FLP recombinase under the indicated promoters. Genomic DNA was isolated from whole animals, and PCR was performed using primers that distinguish the intact *smn-1-FRT* allele (“wild-type”) from the recombined deletion allele. Efficient excision is detected only in the presence of the corresponding FLP driver. A table summarizing FLP promoters and their target tissues is shown at right. (B) Kaplan–Meier survival analysis of *smn-1-FRT* animals expressing FLP recombinase in different tissues. No-FLP animals serve as controls. Survival curves illustrate differential effects of tissue-specific *smn-1* excision on organismal lifespan. No significant difference was found between “no FLP” and WT (p = 0.1189). (C) Survival analysis comparing wild-type, *smn-1(ok355)* null mutants, and *smn-1(ok355)* animals rescued by intestinal expression of smn-1(ok355) (*smn-1(ok355) ; Ex[Pelt-2::smn-1])*. Intestinal expression of *SMN-1* partially restores survival relative to null mutants. (D) Pharyngeal pumping rates measured across developmental stages (34, 48, 72 and 96 hours post-hatching) in wild-type, *smn-1(ok355)* null mutants, and intestinal rescue animals (*smn-1(ok355); Ex[pelt-2::SMN-1]*). Intestinal expression of *SMN-1* partially restores pumping defects observed in null mutants. Bars represent mean ± SEM. Statistical significance was assessed using one-way ANOVA with appropriate multiple comparisons; ****p < 0.0001.

**Figure S2:**
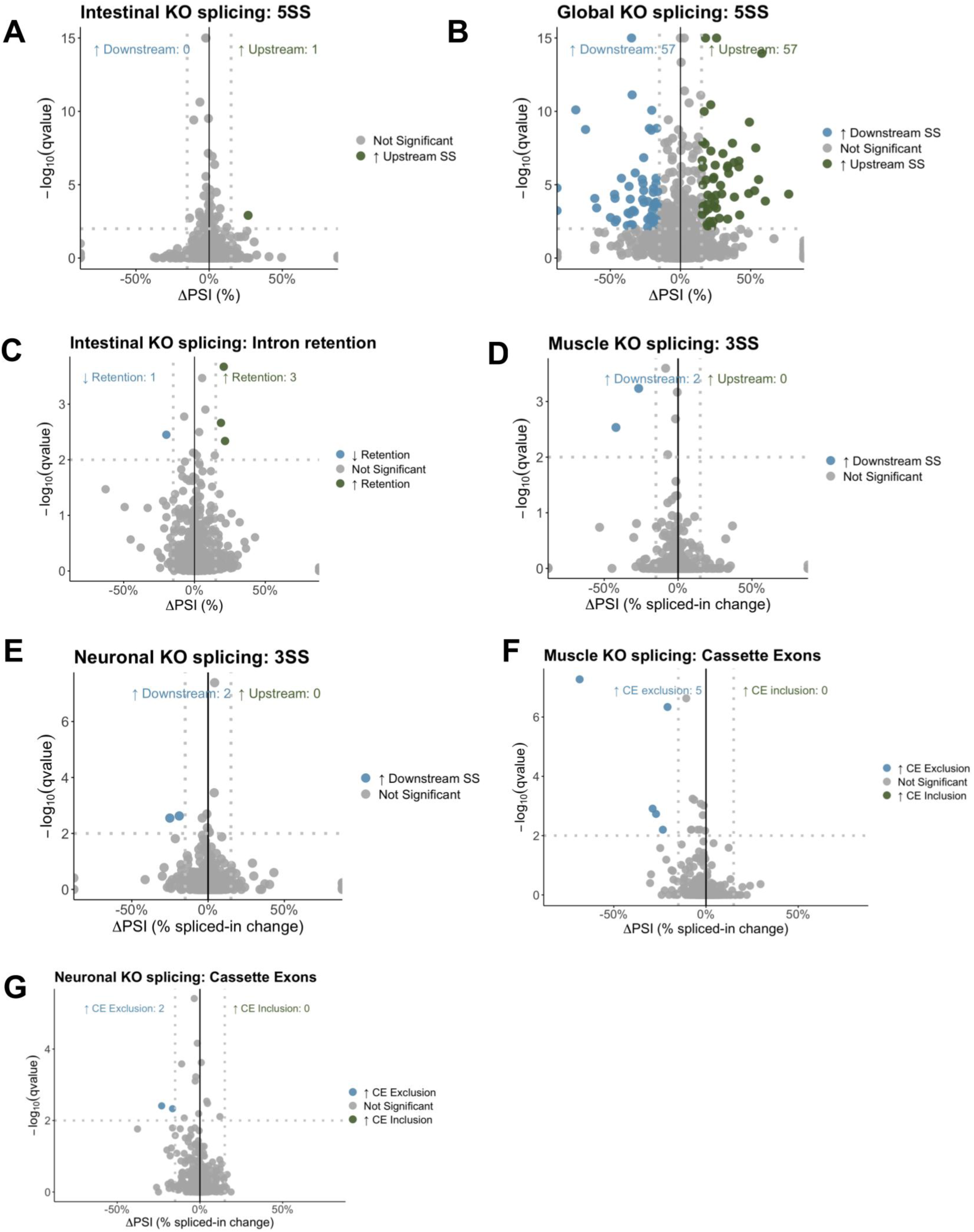
Changes in alternative splicing upon tissue-specific loss of *smn-1*. (A-G) Volcano plot showing differential alternative 5′ splice site (5′SS) usage in intestinal (A) and global (B) *smn-1* knockout animals, intron retention events in intestinal KOs (C), 3’ splice site (3’SS) usage in muscle (D) and neuronal (E) KOs, and cassette exon events in muscle (F) and neuronal (G) KO animals compared to wild-type controls. The x-axis represents the change in percent spliced in (ΔPSI), and the y-axis shows −log10(q value). Events with significantly increased usage of upstream 5′ splice sites are shown in green, events with increased usage of downstream 5′ splice sites are shown in blue, and non-significant events are shown in gray. Dashed vertical lines indicate the ΔPSI cutoff (PSI ≥ 15%), and the horizontal dashed line denotes the significance threshold (q ≤ 0.01).

**Figure S3:**
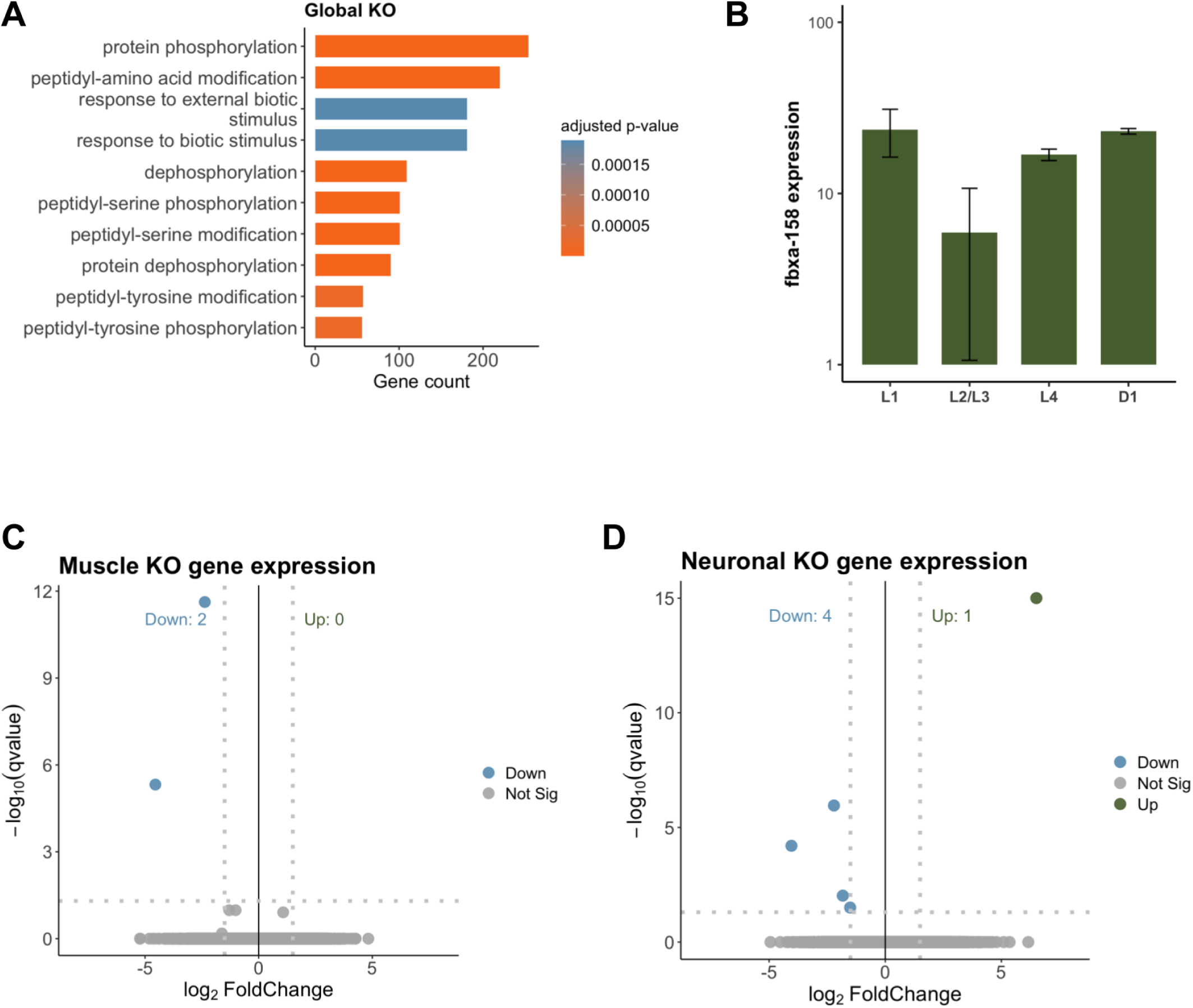
Transcriptional consequences of *smn-1* loss. (A) Gene Ontology (GO) enrichment analysis of differentially expressed genes in global *smn-1* knockout animals. Enriched biological process terms are shown, with bar length indicating the number of genes associated with each term and color representing the adjusted p-value. In addition to immune-related pathways, global loss of *smn-1* is associated with significant enrichment of protein phosphorylation and post-translational modification–related processes. (B) Developmental expression analysis of *fbxa-158* in intestinal *smn-1* knockout animals across larval stages (13,24,34 and 48 hours post-hatching). Expression values are shown on a logarithmic scale and represent mean ± SEM from biological replicates. Elevated *fbxa-158* expression is detected early in development, preceding the onset of overt phenotypic defects. (C, D) Volcano plots of differentially expressed genes in muscle (C) and neuronal KO (D) animals. Log2 fold change is plotted against −log10 adjusted p-value, with significantly upregulated (green) and downregulated (blue) genes indicated.

## REFERENCES

1. Lorson C, Hahnen E, Androphy E, Wirth B. A single nucleotide in the SMN gene regulates splicing and is responsible for spinal muscular atrophy. Proc Natl Acad Sci. 1999;96(11):6307–6311. doi:10.1073/pnas.96.11.6307

2. Liu Q, Dreyfuss G. A novel nuclear structure containing the survival of motor neurons protein. EMBO J. 1996;15(14):3555–3565. doi:10.1002/j.1460-2075.1996.tb00725.x

3. Monani UR. Spinal Muscular Atrophy: A Deficiency in a Ubiquitous Protein; a Motor Neuron-Specific Disease. Neuron. 2005;48(6):885–895. doi:10.1016/j.neuron.2005.12.001

4. Lefebvre S, Bürglen L, Reboullet S, et al. Identification and characterization of a spinal muscular atrophy-determining gene. Cell. 1995;80(1):155–165. doi:10.1016/0092-8674(95)90460-3

5. Vitali T, Sossi V, Tiziano F, et al. Detection of the Survival Motor Neuron (SMN) Genes by FISH: Further Evidence for a Role for SMN2 in the Modulation of Disease Severity in SMA Patients. Hum Mol Genet. 1999;8(13):2525–2532. doi:10.1093/hmg/8.13.2525

6. Singh NK, Singh NN, Androphy EJ, Singh RN. Splicing of a Critical Exon of Human Survival Motor Neuron Is Regulated by a Unique Silencer Element Located in the Last Intron. Mol Cell Biol. 2006;26(4):1333–1346. doi:10.1128/MCB.26.4.1333-1346.2006

7. Chaytow H, Huang YT, Gillingwater TH, Faller KME. The role of survival motor neuron protein (SMN) in protein homeostasis. Cell Mol Life Sci CMLS. 2018;75(21):3877–3894. doi:10.1007/s00018-018-2849-1

8. Faravelli I, Riboldi GM, Rinchetti P, Lotti F. The SMN Complex at the Crossroad between RNA Metabolism and Neurodegeneration. Int J Mol Sci. 2023;24(3):2247. doi:10.3390/ijms24032247

9. Gubitz AK, Feng W, Dreyfuss G. The SMN complex. Exp Cell Res. 2004;296(1):51–56. doi:10.1016/j.yexcr.2004.03.022

10. Sintusek P, Catapano F, Angkathunkayul N, et al. Histopathological Defects in Intestine in Severe Spinal Muscular Atrophy Mice Are Improved by Systemic Antisense Oligonucleotide Treatment. PLoS ONE. 2016;11(5):e0155032. doi:10.1371/journal.pone.0155032

11. Gombash SE, Cowley CJ, Fitzgerald JA, et al. SMN deficiency disrupts gastrointestinal and enteric nervous system function in mice. Hum Mol Genet. 2015;24(13):3847–3860. doi:10.1093/hmg/ddv127

12. Shababi M, Lorson CL, Rudnik-Schöneborn SS. Spinal muscular atrophy: a motor neuron disorder or a multi-organ disease? J Anat. 2014;224(1):15–28. doi:10.1111/joa.12083

13. Kolb SJ, Battle DJ, Dreyfuss G. Molecular Functions of the SMN Complex. J Child Neurol. 2007;22(8):990–994. doi:10.1177/0883073807305666

14. Thelen MP, Kye MJ. The Role of RNA Binding Proteins for Local mRNA Translation: Implications in Neurological Disorders. Front Mol Biosci. 2020;Volume 6-2019. https://www.frontiersin.org/journals/molecular-biosciences/articles/10.3389/fmolb.2019.00161

15. Burghes AHM, Beattie CE. Spinal Muscular Atrophy: Why do low levels of SMN make motor neurons sick? Nat Rev Neurosci. 2009;10(8):597–609. doi:10.1038/nrn2670

16. Gallotta I, Mazzarella N, Donato A, et al. Neuron-specific knock-down of SMN1 causes neuron degeneration and death through an apoptotic mechanism. Hum Mol Genet. 2016;25(12):2564–2577. doi:10.1093/hmg/ddw119

17. Dimitriadi M, Derdowski A, Kalloo G, et al. Decreased function of survival motor neuron protein impairs endocytic pathways. Proc Natl Acad Sci. 2016;113(30). doi:10.1073/pnas.1600015113

18. Briese M, Esmaeili B, Fraboulet S, et al. Deletion of smn-1, the Caenorhabditis elegans ortholog of the spinal muscular atrophy gene, results in locomotor dysfunction and reduced lifespan. Hum Mol Genet. 2008;18(1):97–104. doi:10.1093/hmg/ddn320

19. Miguel-Aliaga I, Culetto E, Walker DS, Baylis HA, Sattelle DB, Davies KE. The Caenorhabditis Elegans Orthologue of the Human Gene Responsible for Spinal Muscular Atrophy Is a Maternal Product Critical for Germline Maturation and Embryonic Viability. Hum Mol Genet. 1999;8(12):2133–2143. doi:10.1093/hmg/8.12.2133

20. Gao X, Teng Y, Luo J, et al. The survival motor neuron gene smn-1 interacts with the U2AF large subunit gene uaf-1 to regulate Caenorhabditis elegans lifespan and motor functions. RNA Biol. 2014;11(9):1148–1160. doi:10.4161/rna.36100

21. Coovert DD, Le TT, McAndrew PE, et al. The Survival Motor Neuron Protein in Spinal Muscular Atrophy. Hum Mol Genet. 1997;6(8):1205–1214. doi:10.1093/hmg/6.8.1205

22. Burlet P, Huber C, Bertrandy S, et al. The distribution of SMN protein complex in human fetal tissues and its alteration in spinal muscular atrophy. Hum Mol Genet. 1998;7(12):1927–1933. doi:10.1093/hmg/7.12.1927

23. Gonzalez D, Vásquez-Doorman C, Luna A, Allende ML. Modeling Spinal Muscular Atrophy in Zebrafish: Current Advances and Future Perspectives. Int J Mol Sci. 2024;25(4):1962. doi:10.3390/ijms25041962

24. Groen EJN, Perenthaler E, Courtney NL, et al. Temporal and tissue-specific variability of SMN protein levels in mouse models of spinal muscular atrophy. Hum Mol Genet. 2018;27(16):2851–2862. doi:10.1093/hmg/ddy195

25. Muñoz-Jiménez C, Ayuso C, Dobrzynska A, Torres-Mendéz A, Ruiz PDLC, Askjaer P. An Efficient FLP-Based Toolkit for Spatiotemporal Control of Gene Expression in *Caenorhabditis elegans*. Genetics. 2017;206(4):1763–1778. doi:10.1534/genetics.117.201012

26. Macías-León J, Askjaer P. Efficient FLP-mediated germ-line recombination in C. elegans. MicroPublication Biol. 2018:10.17912/W2G66S. doi:10.17912/W2G66S

27. Trojanowski NF, Raizen DM, Fang-Yen C. Pharyngeal pumping in Caenorhabditis elegans depends on tonic and phasic signaling from the nervous system. Sci Rep. 2016;6(1):22940. doi:10.1038/srep22940

28. Avery L, You YJ. C. elegans feeding. In: WormBook: The Online Review of C. Elegans Biology [Internet]. WormBook; 2018. Accessed September 18, 2025. https://www.ncbi.nlm.nih.gov/books/NBK116080/

29. Dambroise E, Monnier L, Ruisheng L, et al. Two phases of aging separated by the Smurf transition as a public path to death. Sci Rep. 2016;6:23523. doi:10.1038/srep23523

30. Liu D, Thomas J. Regulation of a periodic motor program in C. elegans. J Neurosci. 1994;14(4):1953–1962. doi:10.1523/JNEUROSCI.14-04-01953.1994

31. Gao X, Xu J, Chen H, et al. Defective Expression of Mitochondrial, Vacuolar H+-ATPase and Histone Genes in a C. elegans Model of SMA. Front Genet. 2019;10. doi:10.3389/fgene.2019.00410

32. Osman EY, Bolding MR, Villalón E, et al. Functional characterization of SMN evolution in mouse models of SMA. Sci Rep. 2019;9(1):9472. doi:10.1038/s41598-019-45822-8

33. Fischer U, Liu Q, Dreyfuss G. The SMN–SIP1 Complex Has an Essential Role in Spliceosomal snRNP Biogenesis. Cell. 1997;90(6):1023–1029. doi:10.1016/S0092-8674(00)80368-2

34. Tsuiji H, Iguchi Y, Furuya A, et al. Spliceosome integrity is defective in the motor neuron diseases ALS and SMA. EMBO Mol Med. 2013;5(2):221–234. doi:10.1002/emmm.201202303

35. Rizzo F, Nizzardo M, Vashisht S, et al. Key role of SMN/SYNCRIP and RNA-Motif 7 in spinal muscular atrophy: RNA-Seq and motif analysis of human motor neurons. Brain. 2019;142(2):276–294. doi:10.1093/brain/awy330

36. Rashid S, Shen A, Yong A, Akay A, Dimitriadi M. Widespread intron retention and exon skipping characterise alternative splicing changes in a C. elegans model of spinal muscular atrophy. Hum Mol Genet. Published online December 1, 2025:ddaf176. doi:10.1093/hmg/ddaf176

37. Reddy KC, Dror T, Sowa JN, et al. An Intracellular Pathogen Response Pathway Promotes Proteostasis in C. elegans. Curr Biol. 2017;27(22):3544–3553.e5. doi:10.1016/j.cub.2017.10.009

38. Panek J, Gang SS, Reddy KC, et al. A cullin-RING ubiquitin ligase promotes thermotolerance as part of the intracellular pathogen response in *Caenorhabditis elegans*. Proc Natl Acad Sci. 2020;117(14):7950–7960. doi:10.1073/pnas.1918417117

39. Lažetić V, Wu F, Cohen LB, et al. The transcription factor ZIP-1 promotes resistance to intracellular infection in Caenorhabditis elegans. Nat Commun. 2022;13(1):17. doi:10.1038/s41467-021-27621-w

40. Lažetić V, Blanchard MJ, Bui T, Troemel ER. Multiple pals gene modules control a balance between immunity and development in Caenorhabditis elegans. bioRxiv. Published online January 18, 2023:2023.01.15.524171. doi:10.1101/2023.01.15.524171

41. Lažetić V, Batachari LE, Russell AB, Troemel ER. Similarities in the induction of the intracellular pathogen response in Caenorhabditis elegans and the type I interferon response in mammals. Bioessays. 2023;45(11):2300097. doi:10.1002/bies.202300097

42. Leyva-Díaz E, Stefanakis N, Carrera I, et al. Silencing of Repetitive DNA Is Controlled by a Member of an Unusual *Caenorhabditis elegans* Gene Family. Genetics. 2017;207(2):529–545. doi:10.1534/genetics.117.300134

43. Ji L, Wang Y, Zhou L, et al. E3 Ubiquitin Ligases: The Operators of the Ubiquitin Code That Regulates the RLR and cGAS-STING Pathways. Int J Mol Sci. 2022;23(23):14601. doi:10.3390/ijms232314601

44. Garcia-Sanchez JA, Ewbank JJ, Visvikis O. Ubiquitin-related processes and innate immunity in C. elegans. Cell Mol Life Sci CMLS. 2021;78(9):4305–4333. doi:10.1007/s00018-021-03787-w

45. Bakowski MA, Desjardins CA, Smelkinson MG, et al. Ubiquitin-Mediated Response to Microsporidia and Virus Infection in C. elegans. Schneider DS, ed. PLoS Pathog. 2014;10(6):e1004200. doi:10.1371/journal.ppat.1004200

46. Kuhn DJ, Chen Q, Voorhees PM, et al. Potent activity of carfilzomib, a novel, irreversible inhibitor of the ubiquitin-proteasome pathway, against preclinical models of multiple myeloma. Blood. 2007;110(9):3281–3290. doi:10.1182/blood-2007-01-065888

47. Kupperman E, Lee EC, Cao Y, et al. Evaluation of the proteasome inhibitor MLN9708 in preclinical models of human cancer. Cancer Res. 2010;70(5):1970–1980. doi:10.1158/0008-5472.CAN-09-2766

48. Anderson DJ, Le Moigne R, Djakovic S, et al. Targeting the AAA ATPase p97 as an Approach to Treat Cancer through Disruption of Protein Homeostasis. Cancer Cell. 2015;28(5):653–665. doi:10.1016/j.ccell.2015.10.002

49. Ghaddar A, Armingol E, Huynh C, et al. Whole-body gene expression atlas of an adult metazoan. Sci Adv. 9(25):eadg0506. doi:10.1126/sciadv.adg0506

50. Wolfe Z, Liska D, Norris A. Deep transcriptomics reveals cell-specific isoforms of pan-neuronal genes. Nat Commun. 2025;16:4507. doi:10.1038/s41467-025-58296-2

51. Gogliotti RG, Quinlan KA, Barlow CB, Heier CR, Heckman CJ, DiDonato CJ. Motor Neuron Rescue in Spinal Muscular Atrophy Mice Demonstrates That Sensory-Motor Defects Are a Consequence, Not a Cause, of Motor Neuron Dysfunction. J Neurosci. 2012;32(11):3818–3829. doi:10.1523/JNEUROSCI.5775-11.2012

52. Park GH, Maeno-Hikichi Y, Awano T, Landmesser LT, Monani UR. Reduced Survival of Motor Neuron (SMN) Protein in Motor Neuronal Progenitors Functions Cell Autonomously to Cause Spinal Muscular Atrophy in Model Mice Expressing the Human Centromeric (SMN2) Gene. J Neurosci. 2010;30(36):12005–12019. doi:10.1523/JNEUROSCI.2208-10.2010

53. Jablonka S, Schrank B, Kralewski M, Rossoll W, Sendtner M. Reduced survival motor neuron (Smn) gene dose in mice leads to motor neuron degeneration: an animal model for spinal muscular atrophy type III. Hum Mol Genet. 2000;9(3):341–346. doi:10.1093/hmg/9.3.341

54. Gallotta I, Mazzarella N, Donato A, et al. Neuron-specific knock-down of SMN1 causes neuron degeneration and death through an apoptotic mechanism. Hum Mol Genet. 2016;25(12):2564–2577. doi:10.1093/hmg/ddw119

55. Walsh MB, Janzen E, Wingrove E, et al. Genetic modifiers ameliorate endocytic and neuromuscular defects in a model of spinal muscular atrophy. BMC Biol. 2020;18(1):127. doi:10.1186/s12915-020-00845-w

56. Sleigh JN, Buckingham SD, Esmaeili B, et al. A novel Caenorhabditis elegans allele, smn-1(cb131), mimicking a mild form of spinal muscular atrophy, provides a convenient drug screening platform highlighting new and pre-approved compounds. Hum Mol Genet. 2011;20(2):245–260. doi:10.1093/hmg/ddq459

57. Silva-García CG, Lanjuin A, Heintz C, Dutta S, Clark NM, Mair WB. Single-Copy Knock-In Loci for Defined Gene Expression in Caenorhabditis elegans. G3 GenesGenomesGenetics. 2019;9(7):2195–2198. doi:10.1534/g3.119.400314

58. Yeo CJJ, Darras BT. Overturning the Paradigm of Spinal Muscular Atrophy as Just a Motor Neuron Disease. Pediatr Neurol. 2020;109:12–19. doi:10.1016/j.pediatrneurol.2020.01.003

59. Asakawa K, Tomita T, Shioya S, Handa H, Saeki Y, Kawakami K. Intrinsically accelerated cellular degradation is amplified by TDP-43 loss in ALS-vulnerable motor neurons in a zebrafish model. Nat Commun. 2025;16(1):9213. doi:10.1038/s41467-025-65097-0

60. Ng SY, Soh BS, Rodriguez-Muela N, et al. Genome-Wide RNA-Seq of Human Motor Neurons Implicates Selective ER Stress Activation in Spinal Muscular Atrophy. Cell Stem Cell. 2015;17(5):569–584. doi:10.1016/j.stem.2015.08.003

61. Sowa JN, Jiang H, Somasundaram L, et al. The Caenorhabditis elegans RIG-I Homolog DRH-1 Mediates the Intracellular Pathogen Response upon Viral Infection. J Virol. 2020;94(2):e01173–19. doi:10.1128/JVI.01173-19

62. Lažetić V, Batachari LE, Russell AB, Troemel ER. Similarities in the induction of the intracellular pathogen response in Caenorhabditis elegans and the type I interferon response in mammals. Bioessays. 2023;45(11):2300097. doi:10.1002/bies.202300097

63. Garcia EL, Steiner RE, Raimer AC, Herring LE, Matera AG, Spring AM. Dysregulation of innate immune signaling in animal models of spinal muscular atrophy. BMC Biol. 2024;22:94. doi:10.1186/s12915-024-01888-z

64. Deguise MO, Kothary R. New insights into SMA pathogenesis: immune dysfunction and neuroinflammation. Ann Clin Transl Neurol. 2017;4(7):522–530. doi:10.1002/acn3.423

65. Kwon DY, Motley WW, Fischbeck KH, Burnett BG. Increasing expression and decreasing degradation of SMN ameliorate the spinal muscular atrophy phenotype in mice. Hum Mol Genet. 2011;20(18):3667–3677. doi:10.1093/hmg/ddr288

66. Brenner S. The Genetics of CAENORHABDITIS ELEGANS. Genetics. 1974;77(1):71–94. doi:10.1093/genetics/77.1.71

67. Large-Scale Screening for Targeted Knockouts in the Caenorhabditis elegans Genome. G3 GenesGenomesGenetics. 2012;2(11):1415–1425. doi:10.1534/g3.112.003830

68. Sleigh JN, Gillingwater TH, Talbot K. The contribution of mouse models to understanding the pathogenesis of spinal muscular atrophy. Dis Model Mech. 2011;4(4):457–467. doi:10.1242/dmm.007245

69. Napier-Jameson R, Marx O, Norris A. A pair of RNA binding proteins inhibit ion transporter expression to maintain lifespan. Genetics. 2023;226(2):iyad212. doi:10.1093/genetics/iyad212

70. Dobin A, Davis CA, Schlesinger F, et al. STAR: ultrafast universal RNA-Seq aligner. Bioinformatics. 2013;29(1):15–21. doi:10.1093/bioinformatics/bts635

71. Love MI, Huber W, Anders S. Moderated estimation of fold change and dispersion for RNA-Seq data with DESeq2. Genome Biol. 2014;15(12):550. doi:10.1186/s13059-014-0550-8

72. Wang Q, Rio DC. JUM is a computational method for comprehensive annotation-free analysis of alternative pre-mRNA splicing patterns. Proc Natl Acad Sci U S A. 2018;115(35):E8181–E8190. doi:10.1073/pnas.1806018115

